# An *ex vivo* and *in vitro* investigation of extracellular vesicle interactions with B cells of *Macaca nemestrina* and humans

**DOI:** 10.1101/2025.02.12.637883

**Authors:** Bianca C. Pachane, Blanca V. Rodriguez, Erin N. Shirk, Olesia Gololobova, Bess Carlson, Suzanne E. Queen, Loren D. Erickson, Heloisa S. Selistre-de-Araujo, Kenneth W. Witwer

## Abstract

Extracellular vesicles may modify recipient cell behavior through multiple mechanisms, including interacting with the cell surface or internal membrane components and delivering luminal cargo to the cytoplasm. Here, we use a previously established *ex vivo* approach to investigate the cellular fate of EVs spiked into whole blood samples from nonhuman primate (NHP) and human donors and contrast these findings with results from *in vitro* assays. We report that EVs are internalized by NHP and human B cells while also associating to some degree with other PBMCs. EVs exhibit greater association with B cells in *ex vivo* whole blood compared to isolated B cells, suggesting that blood components may promote EV interactions or that cell isolation factors may inhibit this association. Cellular uptake of EVs involves clathrin-dependent endocytosis and may be aided by other pathways, including direct EV-cell membrane fusion. Overall, our data suggest that EV association, including uptake, by B cells occurs in at least two primate species. These findings highlight the potential to develop new strategies to either enhance or inhibit EV tropism toward B cells.

**Graphical Abstract:** 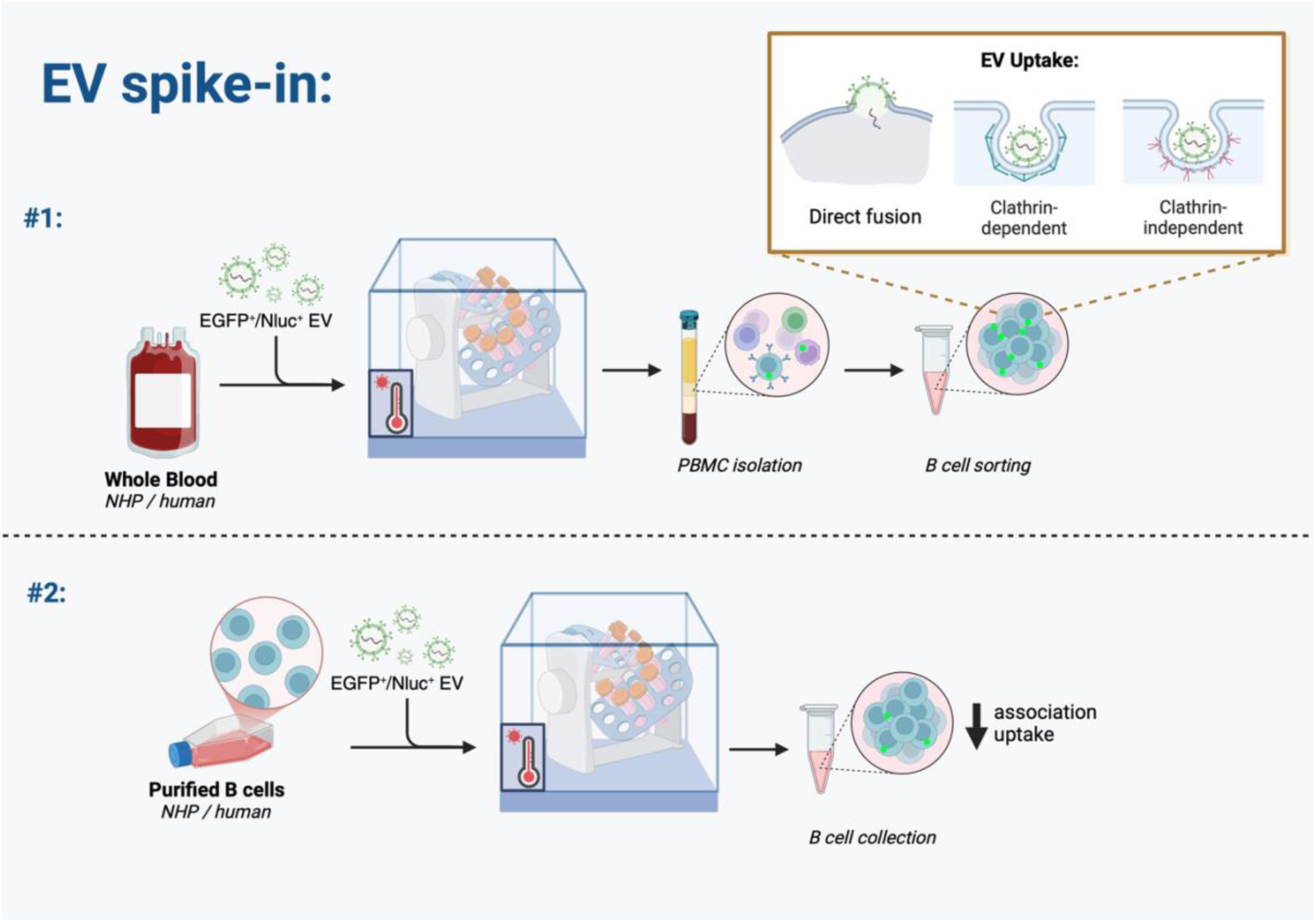

## Introduction

Multiple cell behaviors can be modulated by extracellular vesicles (EVs), which can interact with the cell surface and/or become internalized (1) through multi-step processes (2). An EV that encounters a cell may first bind to the surface via adhesion proteins including tetraspanins and integrins. The EV may then remain attached to the surface, detach and move away, or be internalized into the cell, for example, by clathrin-dependent mechanisms, clathrin-independent pathways, or direct membrane fusion (1,2). If the EV fuses directly with the plasma membrane, or if uptake is followed by EV-cell membrane fusion, vesicular cargo is released into the cytoplasm. Alternatively, in the absence of EV fusion, the EV may be degraded in the lysosome or released back into the extracellular space via endosome-plasma membrane fusion (3).

EV internalization into recipient cells is thought to occur mainly through clathrin-mediated endocytosis, whereby a scaffold comprised of heavy and light clathrin chains is assembled around an endocytic vesicle, enabling actin-mediated direction to endosomal or Golgi pathways (4). Clathrin-independent mechanisms include caveolin-mediated endocytosis, a process in which CAV1 binds to lipid microdomains on the cell membrane to effect internalization (2,5). By this mechanism, the vesicle is fused with the early endosome for recycling or associated with multivesicular bodies for ubiquitination (5). Small EV internalization can also occur through macropinocytosis, a mechanism triggered by PIP3 signaling that promotes actin polymerization and enables the cell membrane to expand and encapsulate the region of interest (2,3,6).

In this study, we sought to expand our knowledge of EV interactions with B cells, an apparent, previously underappreciated target of some EVs in the primate bloodstream (7). While most studies of EV biodistribution have understandably focused on murine models, using a combination of *in vivo, in situ*, and *ex vivo* techniques (8,9), we recently observed strong interactions between intravenously administered EVs and B cells of NHP (7). Similar results were found in an *ex vivo* model in which NHP donor blood was drawn and spiked with exogenous EVs (10). In this study, we use this *ex vivo* model to further investigate the interactions of EVs with B cells from both NHP and human donors, including EV uptake mechanisms.

## Materials and Methods

### EV production

EVs were separated from Expi293F cells transiently transfected with pLenti-palmGRET, as previously described (7,10,11). Briefly, the cell suspension was collected three days after transfection and spun twice to remove cells and debris (300 × g for 5 min; 2,000 × g for 20 min). The supernatant was filtered through a 0.22 µm bottle-top system (Corning), concentrated 10X by tangential flow filtration (TFF, Vivaflow^®^ 50R TFF cassettes, Sartorius) and 4X by ultrafiltration (Centricon Plus 70 Ultracel® PL-100, Merck Millipore; 4,000 × g, 20 min, RT). After elution, samples were separated using size-exclusion chromatography (SEC) into 15 fractions using qEV10 70nm columns (iZON). EV-enriched fractions (1–4) were pooled and concentrated using Amicon 15 Ultra RC - 10 kDa MWCO filters (Merck Millipore). Expi293F-palmGRET EV aliquots in Dulbecco’s phosphate-buffered saline (DPBS) were stored at 4 °C for short-term use and -80 °C for long-term storage. We have submitted all relevant EV separation and characterization data to the EV-TRACK knowledgebase (EV-TRACK ID: EV250009) (12). EV separation and characterization were conducted following the MISEV guidelines (13,14).

### Western blotting

Cell lysates were prepared from an Expi293F-palmGRET cell suspension (3 x 10^6^ cells/ml) harvested from culture, washed with DPBS (2,500 × g, 5 min, RT) and mixed with RIPA buffer (Cell Signaling Technology #9806) for 1 h on ice. The supernatant was collected after centrifugation (14,000 × g for 15 min at 4 °C) and quantified using BCA protein assay (Thermo Scientific #23227). Expi293F-palmGRET EVs were lysed with RIPA buffer for 10 min at room temperature. Samples (10 µl) were mixed with sample buffer (reducing: Thermo Scientific #39000, or non-reducing: Thermo Scientific #39001), boiled at 100 °C for 5 min, and loaded into Criterion TGX stain-free gels (4-15% pre-cast, 18 wells; 5678084, Bio-Rad) with Spectra™ Multicolor Broad Range Protein Ladder (26634, Thermo Fisher) as a loading control. Gels ran in Tris/Glycine/SDS buffer (1610772, Bio-Rad) at constant voltage (100V) for 1.5 h, and stain-free images were obtained on a Gel Doc EZ imager (Bio-Rad). Proteins were transferred to PVDF membranes (iBlot™ 2 Transfer Stacks, Invitrogen #IB24001) using a stacked program (20 V for 1 min; 23 V for 4 min, and 25 V for 2 min) from the iBlot™ 2 Gel Transfer Device (Invitrogen). Under agitation, membranes were blocked for 1 h in 5% milk-PBST and exposed to primary antibodies overnight at 4 °C. For EV characterization, membranes were probed for CD63 (1:3000, Ms, BD #556016), CD9 (1:3000, Ms, BioLegend #312102), syntenin (1:1000, Rb, Abcam #133267), Alix (1:1000, Rb, Abcam #186429) and calnexin (1:1000, Rb, Abcam #22595). For cell fractionation, membranes were probed for β-actin (1:1000, Cell Signaling #3700), calnexin (1:1000, Abcam #22595), and GAPDH (1:2500, Abcam #9485). Following four 5-minute washes with 5% milk-PBST, membranes were incubated with the appropriate secondary antibodies (m-IgGκ BP-HRP, 1:5000, Santa Cruz #616102; or Mouse anti-Rb IgG HRP, 1:5000, Santa Cruz #2327) for 1h under agitation at room temperature. Membranes were washed three times with PBST and exposed for 60 seconds to SuperSignal™ West Pico ECL substrate (Thermo Scientific #34578) before imaging on iBright 1500 (Invitrogen). Integer stain-free gels and membranes are available in the Supporting Information section (Suppl. Fig. 1).

### Single-particle interferometric reflectance imaging sensing

Chips from the human tetraspanin kit (Ref EV-TETRA-C, Lot NV221653001D, kit version EV-RGT-02, Unchained Labs) were pre-scanned using an ExoView reader and Exoview Analyzer software before the start of the assay. Expi293F-palmGRET EVs (4E+8 particles/ml) in 35 µl incubation solution were incubated on top of chips overnight at RT, protected from light. Using the CW100 chip washer and the program CW-TETRA, chips were washed and incubated with CD63 and CD81 antibodies following the manufacturer’s recommendations. Chips were moved to the 45° ramp for draining and transferred to an absorbent pad to dry completely before being loaded into the ExoView dock for analysis.

### NanoFlow cytometry

Measurements of particle concentration, size distribution, and EGFP positivity (%) of EV samples were obtained using an NFCM Flow NanoAnalyzer (NanoFCM Co., Ltd). The system was pre-calibrated for particle concentration with fluorescent 250 nm silica nanoparticles (2.19E+10 particles/ml, NanoFCM #QS2503) and, for sizing, with a premixed silica nanosphere cocktail with monodisperse nanoparticle populations of 68 nm, 91 nm, 113 nm, and 155 nm in diameter (NanoFCM #516M-Exo). Vehicle (DPBS) was set as the blank. EGFP^+^ EV samples were diluted 1:1,000 (v/v) in DPBS, boosted for 60 seconds, and acquired using constant pressure of 1 kPa and event rates between 1,500 and 10,000 events/min. Side-scattering and EGFP signals, particle concentration and size dispersion were calculated using NanoFCM Professional Suite V2.0 software. All essential information following MIFlowCyt-EV guidelines (15) is available in the Supplementary Information section (Suppl. Inf. 2).

### Transmission electron microscopy

EVs were diluted 1:20 (v/v) and adsorbed to glow-discharged carbon-coated 400 mesh copper grids by flotation for 2 min. Grids were quickly blotted and rinsed by flotation on three drops of Tris-buffered saline, 1 min each, before being negatively stained in two consecutive drops of 1% uranyl acetate (UAT) with tylose (1% UAT in deionized water (diH2O), double filtered through a 0.22-μm filter). Grids were blotted and quickly aspirated to cover the sample with a thin layer of stain. Grids were imaged on a Hitachi 7600 transmission electron microscope (TEM) operating at 80 kV with an AMT XR80 CCD (8 megapixel), under magnification ranging from 30,000x to 120,000x.

### Super-resolution microscopy

Glass coverslips were rinsed with 70% ethanol and placed atop a 30 µl drop of EV dilution (4E+08 particles/ml) on parafilm overnight at 4 °C, protected from light. Coverslips were rinsed twice with DPBS and assembled to histological slides using VECTASHIELD^®^ mounting medium. Slides were sealed (CoverGrip Coverslip Sealant, Biotium #23005) imaged using the super-resolution microscope Nanoimager (ONI), with the NimOS software. The microscope was calibrated with beads excited by 405/473/532/635 nm lasers using a 100x oil-immersion objective. Channel mapping calibration was considered successful if point coverage achieved good quality. EVs were imaged in at least three random sites (79.28 µm x 49.43 µm), using 2000 frames, 30 ms exposure, 1.4 numerical aperture. Laser intensity was optimized for each sample. Post-acquisition spatial and statistical analysis was performed using CODI (alto.codi.bio). Reconstituted images were snapped under 6-8x zoom.

### EV spike-in experiments in whole blood

Non-human primate (NHP) samples were obtained from specific pathogen-free (SPF) juvenile male pigtailed macaques *(Macaca nemestrina)* in procedures approved by the Johns Hopkins University Animal Care and Use Committee (ACUC #PR21M297) and conducted following the Weatherall Report, the Guide for the Care and Use of Laboratory Animals, and the USDA Animal Welfare Act. Whole blood was drawn by venipuncture into sterile Luer-lock syringe tubes (BD #309653) and mixed with acid citrate dextrose (ACD) at a ratio of 5:1 (v/v). Whole blood bags from seven anonymous human volunteer donors were purchased from the New York Blood Center (NYBC, New York, NY). Each blood bag contained approximately 500 ml of whole blood with 70 ml of citrate-phosphate-dextrose (CPD) anticoagulant. Blood samples were processed within one hour of delivery. For time-sensitive experiments, blood was aliquoted into microtubes or conical tubes under sterile conditions and spiked with Expi293F-palmGRET EVs at a concentration of 8E+08 particles/ml based on our previous studies (7,10). Samples were incubated on a rotator in an incubator set at 37 °C, with the “mix” mode selected at a speed of 8 rpm (Benchmark Scientific Roto-Therm Plus Incubated Rotator). EV interaction with whole blood components was monitored at 5, 10, 15, 30, 60, 120, and 240 minutes after spike-in.

### Isolation of PBMCs

NHP blood samples were diluted (1:1, v/v) in Hank’s balanced salt solution (HBSS) and applied to SepMate™-15 (IVD) tubes loaded with Percoll^®^ density gradient medium (Cytiva #17-0891-01). Tubes were centrifuged at 1,200 × g for 10 min for triphasic blood separation, with mononuclear cells (MNC) in the upper fraction. MNCs were transferred to fresh tubes, washed with HBSS, and pelleted at 300 × g for 8 min. Cells were treated with red blood cell lysis buffer (ACK lysing buffer 0.83% NH4Cl, 0.1% KHCO3, 0.03% EDTA) for 10 min at 37 °C, washed with HBSS, and centrifuged at 300 × g for 8 min.

Human donor (HD) blood samples were diluted (1:1, v/v) in washing buffer (PBS + 2% FBS + 2 mM EDTA) and applied to SepMate™-50 (IVD) tubes loaded with Ficoll-Paque™ PLUS density gradient (Cytiva # 17144003). Tubes were centrifuged at 1,200 × g for 10 min for triphasic blood separation, with mononuclear cells (MNC) in the upper fraction. MNCs were transferred to fresh tubes, washed with buffer, and pelleted at 300 × g for 8 min. Cells were treated with red blood cell lysis buffer for 10 min at 37 °C, washed with buffer, and centrifuged at 300 × g for 8 min. PBMC count and viability were assessed using trypan blue exclusion method (Countess II FL, Invitrogen).

### CD20^+^ B cell positive selection

PBMCs were resuspended in cold selection buffer (PBS + 0.5% BSA + 2 mM EDTA) mixed with 20% (v/v) CD20 MicroBeads (Miltenyi Biotec 130-191-104 or #130-191-105) for 15 min at 4 °C. Samples were diluted in selection buffer, spun at 300 × g for 8 min, and resuspended for magnetic cell sorting using MS columns (Miltenyi Biotec #130-042-201) in MiniMACS™ separators (Miltenyi Biotec). CD20^-^ PBMCs and CD20^+^ B cells were collected in conical tubes. After elution, CD20^+^ B cells were spun at 300 × g for 8 min and resuspended in RPMI 1640 + 10% FBS + 1% pen/strep medium before assessing viability and cell count. Purified B cells were cultured in RPMI 1640 + 10% FBS + 1% pen/strep medium for up to two days in 6-well plates at 37 °C, 5% CO2.

### Flow cytometry and trypan blue quenching assay

Expi293F-palmGRET EV interaction with PBMCs was assessed using flow cytometry after 15, 30, and 60 min of spike in whole blood. Isolated NHP and HD PBMCs in staining buffer (BioLegend #420201) were immunolabelled for 20 min at room temperature with an antibody cocktail containing mouse-anti-CD3-V500 (1:30, BD Biosciences #560770), mouse-anti-CD4-PerCP/Cy5.5 (1:7.5, BD Biosciences #552838), mouse-anti-CD8-BV570 (1:60, BioLegend #301038), mouse-anti-CD20-e450 (1:60, Thermo Fisher #48-0209-42), mouse-anti-CD159a-PE (1:30, Beckman Coulter #IM3291U), and mouse-anti-CD14-BV650 (1:30, BioLegend #563419). Labeled PBMCs were washed once with DPBS (300 × g, 8 min), resuspended in DPBS, and aliquoted (500 µl) for PBMC-associated EGFP fluorescence analysis using an LSRFortessa™ flow cytometer (BD Biosciences). PBMC aliquots (500 µl in DPBS) were treated with 0.01% (v/v) of 0.4% trypan blue for EGFP quenching (10,16) and re-run immediately. Fluorescence minus one (FMO) controls for CD159a and CD4 were used to accurately gate GFP, PE, and PerCP/Cy5.5 fluorescence. The gating strategy and essential information following MIFlowCyt guidelines (17) are described in the Supporting Information section (Suppl. Inf. 3-4).

### Immunofluorescence

Suspension cells were attached to poly-L-lysine slides (Electron Microscopy Sciences #63410-02) by centrifugation (1000 rpm, 5 min; Cytospin 2, Shandon) and air dried overnight protected from light. Samples were fixed with 4% paraformaldehyde (PFA-H2O) for 10 min at room temperature and washed twice with DPBS to remove excess. Cell permeabilization was achieved by exposing samples to 0.1% triton X-100 for 5 min at room temperature, followed by two washes with DPBS. Unspecific binding was blocked with 1% BSA-PBS for 1h at 4 °C, and cells were exposed to primary antibodies anti-CD27 (1:500, v/v; Abcam #131254), anti-clathrin heavy chain (1:500, v/v; Cell Signaling #2410S), conjugated antibodies CD20-PE (1:50, v/v; Miltenyi Biotech #130-113-374) and CD14-PE (1:50, v/v; Miltenyi Biotec #130-113-147), and iPhalloidin-647 dye (1:1000, v/v; Abcam #176759) for 16-20h at 4 °C. After washes with DPBS, a secondary antibody (Donkey Anti-Rabbit IgG H&L, Alexa Fluor® 647; 1:1,000, v/v; Abcam #150075) was incubated for 1h at 4 °C on the applicable samples. Excess dye was washed with DPBS before coverslip assembly using Prolong Diamond Antifade Mounting Medium with DAPI (Invitrogen #2465103). The slides were sealed with coverslip sealant (Biotium #23005) and stored at 4 °C until analysis.

### Confocal microscopy

Images were acquired using a Zeiss 880 Airyscan FAST confocal microscope under 40x and 63x magnification, using the Zen 2.3 SP1 software (Carl Zeiss AG). Channels for DAPI, A488, A568, and A647 (40.3 pinhole, 850 gain master, 5.0 laser intensity) were imaged in Z-stacks, with 0.3 µm intervals using auto Z brightness correction and optimizing sectioning and step. Z-stacks were manually selected based on the apical and basal location of cells. For EGFP pixel intensity analysis, z-stacks were acquired under 63x magnification, using 512 x 512 px frames at speed 7 and averaging 2, with 8-bit depth. Representative images were acquired under 63x magnification, using frames of 1800 x 1800 px at speed 7 and averaging 2, with 16-bit depth. Image handling and quantification of EGFP pixel intensity were executed on FIJI (Image J) (18).

### Nanoluciferase assay

Nanoluciferase activity was measured using the NanoGlo^®^ Luciferase Assay System (N1120, Promega). EV fractions (1:100 dilution (v/v) in PBS), and B cells (1E+05 cells/ml) were seeded in technical duplicates or triplicates in white flat-bottom 96-well plates (Corning #3922). Assay controls included vehicle (i.e., blank) and EV controls (8E+08 particles/ml). Samples were mixed with the Nano-Glo luciferase assay reagent (1:1, v/v) prepared with 50 parts buffer to 1 part furimazine (substrate). Plates were read immediately after reagent addition at 14.6 mm height and with 1000 integration, using SpectraMax iD5 (Molecular Devices).

### Cell fractionation

B cells were collected in PBS + 1% protease inhibitor (PI, Thermo Fisher #161280) and stored at -20 °C. Once thawed, they were spun at 300 × g for 8 min at 4 °C and resuspended in 200 µl PBS + 1% PI. Cells were vortexed for 5 min, passed through a 30 G needle attached to a 1 ml syringe 5 times, and sonicated (3 cycles of 10 s on + 10 s off) for lysis. We spun samples at 1,200 × g for 10 min at 4 °C to remove any remaining intact cells, and at 100,000 × g for 1 h at 4 °C to separate membrane fractions (pellet) from the cytoplasmic fraction (supernatant). Cell fractions were analyzed using a nanoluciferase assay kit. Successful fractionation was confirmed via Western blotting. Full uncut membranes are available in the Supporting Information section (Suppl. Fig. 3).

### Statistical analysis

Raw data were checked for outliers using the ROUT method^7^, assuming a Q = 0.1. Datasets were transformed and/or normalized based on technical controls for each technique. D’Agostino-Pearson omnibus K2 normality test was applied to identify the type of data distribution. Parametric data were analyzed using ANOVA one-way or two-way with Tukey’s multiple comparison test and are displayed in graphs as mean ± SD. Non-parametric data were analyzed using Kruskal-Wallis analysis of variance with Dunn’s multiple comparison test, and are displayed in graphs as median ± interquartile range. *p* values are demonstrated above significant comparison brackets in graphs. Statistical analysis and graphs were executed on GraphPad Prism (v. 10.0.1 (170)).

## Results

### Expi293F-palmGRET EV separation and characterization

Expi293F-palmGRET EVs were produced and characterized as previously described (19) using methods deposited with the EV-TRACK knowledgebase (EV-TRACK ID: 250009) (12). Expi293F transfection with palmGRET was tracked throughout three days (Suppl. Fig. 4A). Cells produced EGFP^+^/Nluc^+^ EVs with typical morphology and strong fluorescence (Suppl. Fig. 4B-C). SP-IRIS assessed EV fluorescence, and the colocalization between labels indicated an enrichment of CD63/CD81 in our samples (Suppl. Fig. 4D-E). Western blotting detected an enrichment of CD9 and EGFP, traces of ALIX, syntenin and CD63, and minimal calnexin incorporation (Suppl. Fig. 4F). Nano-flow estimated that 63.3% of EVs were EGFP^+^ (Suppl. Fig. 4G), with particle diameter ranging from 50 to 120 nm and peaking at 76 nm (Suppl. Fig. 4H). As previously reported, SEC separation was important, as free nanoluciferase eluted in later SEC fractions (6–12) (Suppl. Fig. 4I) (7).

### Interactions of EGFP^+^ EVs with NHP and human CD20^+^ B cells

EGFP^+^/Nluc^+^ EVs were used to investigate the interaction with NHP or human CD20+ B cells upon spike-in into whole blood or mixture with purified B cells, following the diagram shown in Figure 1A-B. In NHP whole blood, the association of EVs with B cells was detected by nanoluciferase as early as 5 minutes after spike-in and reached significant levels (*p* = 0.0009) after 30 minutes (Fig. 1C). A similar association time course was observed in purified NHP B cells, albeit approximately 5-fold weaker in comparison with the whole blood assay (*p* < 0.0001; Fig. 1D), suggesting that components of whole blood may promote EV association with B cells. We confirmed previous reports (7,10) by using flow cytometry and detecting the EGFP signal as early as 10 minutes after spike-in, with 72.53% (±9.57%) positivity of CD20^+^ B cells, which in turn comprise around 5% of NHP PBMCs (*p* < 0.0001; Fig. 1E). Other PBMCs also interacted with EVs, particularly CD14^+^ monocytes (89.6% ± 7.15%), CD14^-/lo^ monocytes (63.68% ± 10.59%), CD4 T lymphocytes (19.63% ± 11.90), CD8 T lymphocytes (6.62% ± 2.76%). Little or no interaction of EVs with CD159a+ NK cells was detected (0.05% ± 0.08%).

**Figure 1.**
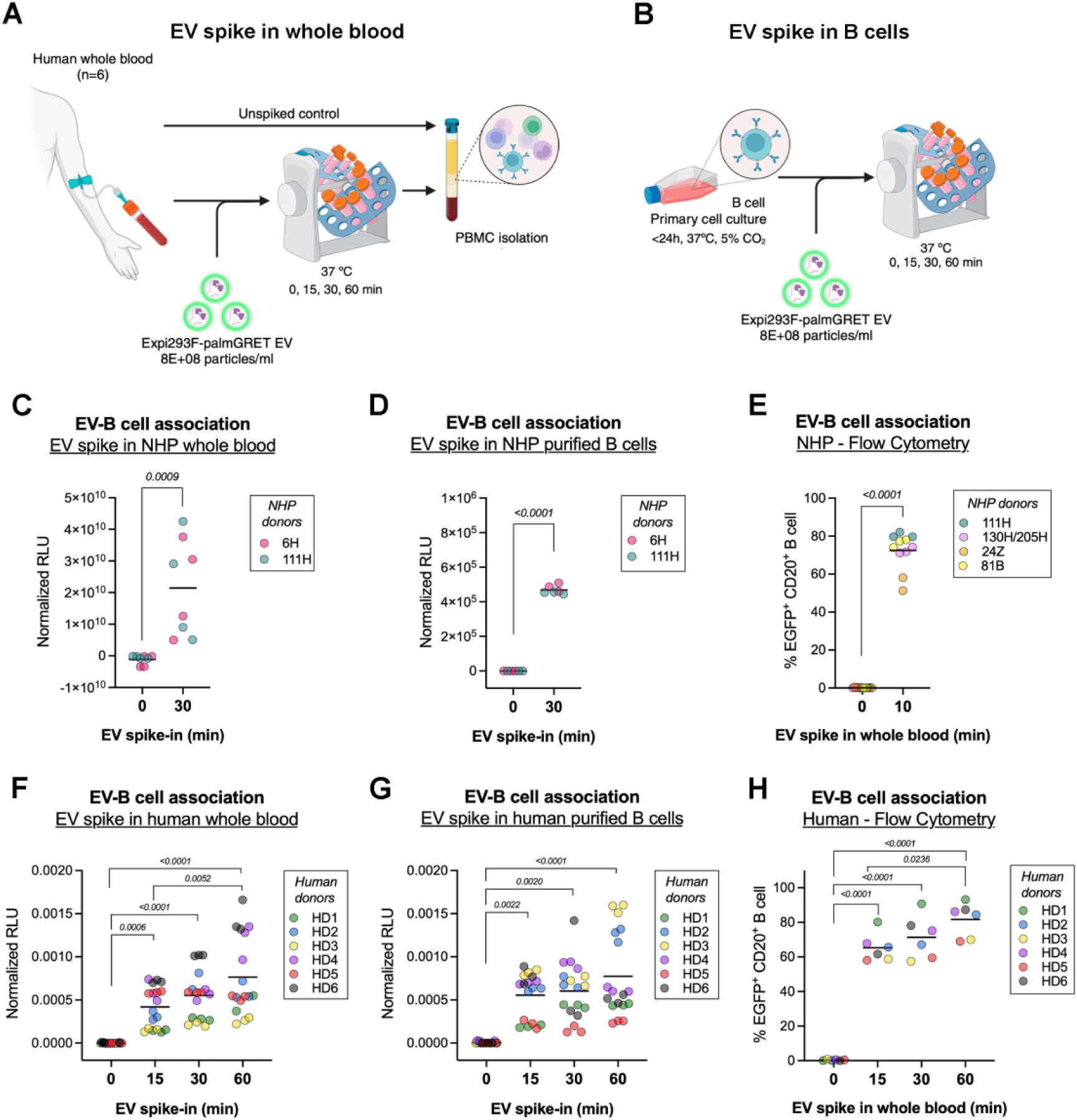
EGFP^+^ EVs interact with NHP and human B cells. (A) Experimental design of EV association with B cells in whole blood. (B) EV association with purified B cells in vitro. Diagrams created using BioRender.com. (C) Nanoluciferase activity of EV association with CD20^+^ B cells after EV spike in NHP whole blood. (D) Nanoluciferase activity of EV association with primary B cells in culture. (E) EGFP^+^ B cells identified by flow cytometry after 10 min of EV spike in NHP whole blood. (F) Nanoluciferase activity of B cells spiked *ex vivo* with EVs. (G) Nanoluciferase activity of B cells spiked *in vitro* with EVs. (H) EGFP^+^ B cells were identified by flow cytometry after 15, 30, and 60 min of EV spike in human whole blood. P values are indicated in brackets.

In human samples, results were similar. Human B cells spiked with EVs in whole blood displayed increasing nanoluciferase signal from 15 minutes (*p* = 0.0006), 30 minutes (*p* < 0.0001), and 60 minutes (*p* <0.0001; Fig. 1F). When purified B cells were exposed to EVs, the relative luminescence followed the same profile (*p* = 0.0022 from 15 min; *p* = 0.0020 from 30 min; *p* < 0.0001from 60 min; Fig. 1G). However, for human samples, the RLU values at each time point were similar for whole blood and purified B cell assays. By flow cytometry, EVs interacted preferentially with CD20^+^ B versus other MNCs (Suppl. Fig. 5A-D). EGFP signal was detected in association with 65.33%, 71.35%, and 81.70% of B cells after 15, 30, and 60 min of EV spike-in into whole blood, respectively (*p* < 0.0001; Fig. 1H).

### NHP and human B cells internalize EGFP^+^ EVs

Using a time-dependent uptake assay (20), we were able to screen EGFP^+^ EV internalization by CD20+ B cells purified from whole blood spike-in experiments. For NHP cells, EGFP signal increased over time, reaching statistically significant levels at 30 minutes (*p* < 0.0001) and increasing until the end of the four-hour assay (*p* < 0.0001; Fig. 2A). Quantification of EGFP signal in human samples indicated a similar response; however, human B cells interacted faster with EVs, leading to a significant rise after 10 minutes of spike-in (*p* < 0.0001; Fig. 2B). Using confocal microscopy to locate EVs within the cell perimeter, we observed the progression of EV uptake by NHP B cells starting at 10 minutes of spike-in, with internalization detected after 30 min and continuing for up to 4 hours (Fig. 2C). As above, human B cells appeared to associate even more quickly, with EVs being detected on the cellular perimeter as early as 5 minutes after spike-in and internalization occurring from 15 minutes onwards (Fig. 2D). Under both experiments, conducted with NHP and human samples, EVs were detected at the cellular surface and internalized, indicating the complexity of interaction between CD20^+^ B cells and EVs. These results also suggest that EV-B cell association and uptake is a conserved mechanism for at least two primate species.

**Figure 2.**
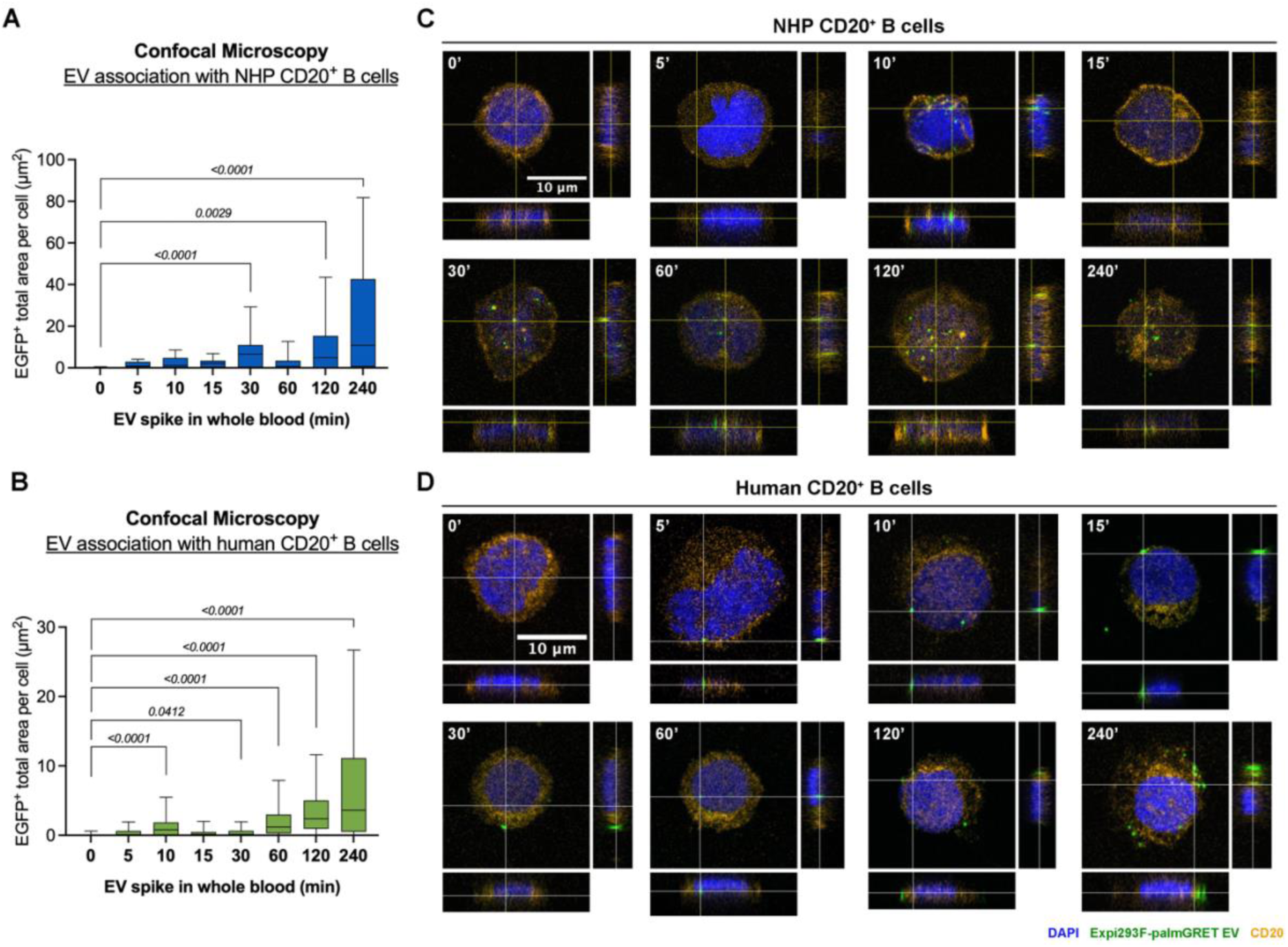
– EV internalization in NHP and human B cells. (A) Quantification of EGFP fluorescence area per NHP B cell over time (n = 80) (B) Representative orthogonal views of NHP B cells in a time-sensitive EV uptake assay. Nuclei stained with DAPI (blue), EGFP-EV (green), and CD20-PE (orange). (C) Quantification of EGFP fluorescence area per human B cell over time (n = 20). (D) Representative orthogonal views of human B cells in a time-sensitive EV uptake assay. Nuclei stained with DAPI (blue), EGFP-EV (green), and CD20-PE (orange). Scale bars: 10 µm. P values are indicated in brackets.

### EV signal distribution in B cells suggests multiple association mechanisms

To assess internalization mechanisms, we first used a flow cytometry uptake assay, in which Trypan blue dye was used to quench “accessible” fluorescent moieties, i.e., those that are not protected inside the cell (16). While 65.33% of human B cells were EGFP^+^ 15 minutes after EV spike-in, only 27.52% had Trypan-inaccessible, or internal, EGFP signal at 15 min (Fig. 3A). EV uptake increased over time, reaching 37.72% of B cells at 30 minutes and 55.33% at one hour. Based on the association and uptake data from flow cytometry, 42.12% of B cells associated with EVs have internalized the particles at 15 minutes of spike-in, increasing to 52.86% at 30 minutes and 65.27% at one hour.

**Figure 3.**
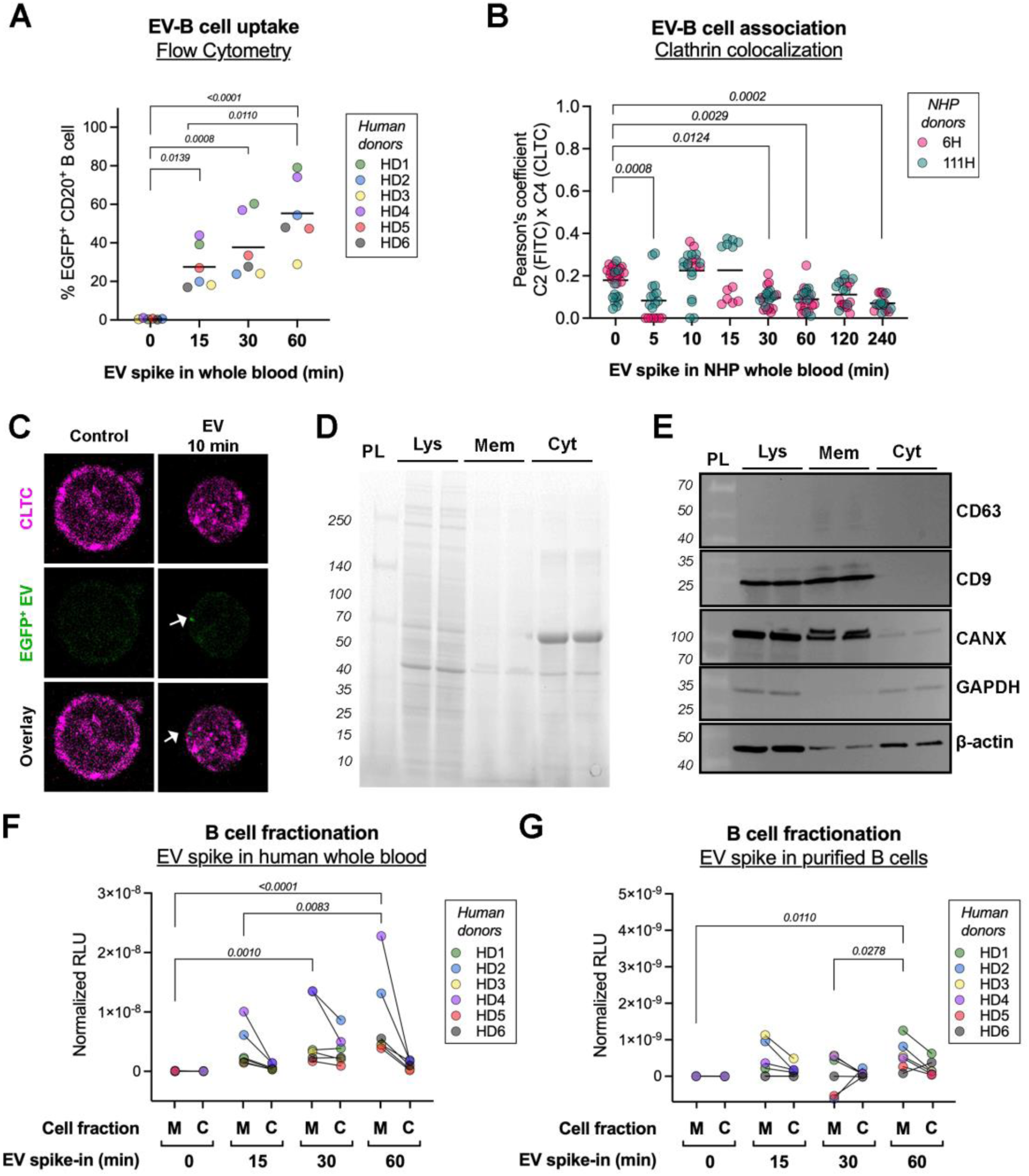
EV signal distribution in B cell compartments. (A) EGFP^+^ B cells quenched with trypan blue, detected by flow cytometry after 15, 30, and 60 min of EV spike in human whole blood. (B) Pearson’s colocalization coefficient values of channels 2 (EGFP) and 4 (clathrin heavy chain, CLTC) from NHP B cells. (C) Representative images of B cells stained for clathrin heavy chain (A647, pink) and Expi293F-palmGRET EVs (EGFP, green). (D) Stain-free gel of B cell fractionation showing different protein profiles between cell lysate (Lys), membrane (Mem) and cytoplasmic (Cyt) fractions. (E) Western blotting membranes probed for CD63, CD9, calnexin, GAPDH and β-actin. PL = protein ladder. (F-G) Analysis of nanoluciferase activity in B cell membrane (M) and cytoplasm (C) fractions after EV spike in (F) human whole blood or (G) purified B cells. P values are indicated in brackets.

As previously discussed, uptake often occurs by clathrin-mediated mechanisms. Since clathrin forms a scaffold around internalized material, we used colocalization analysis to determine if this mechanism is favored. At 5 min of EV spike-in, we observed an initial drop in clathrin heavy chain (CLTC)-EV colocalization, a marker of clathrin involvement, which recovered at 10 and 15 min but declined again at 30 min, remaining stable until the end of the assay (Fig. 3B). The low median values of Pearson’s coefficient, which indicate minimal overlap between signals, suggest that clathrin may facilitate early EV uptake in B cells but is not the sole factor involved in this process. The labeling of CLTC showed its even dispersion around the cell surface throughout all time points (Fig. 3C).

We also examined whether EVs have been dismantled by B cells upon uptake, which may indicate membrane fusion or metabolization of EV content. Human B cells were fractionated to separate membrane and cytoplasmic fractions, with results validated by Western blotting. Each fraction exhibited a distinct protein profile (Fig. 3D), and protein enrichment from each compartment aligned with the expectations. Tetraspanins CD63 and CD9 were found in only lysate and membrane fractions, while GAPDH was exclusive to lysate and cytoplasmic fractions. Two compartment control proteins were found in all samples but more abundantly in one fraction: calnexin, a protein from the endoplasmic reticulum, was found mostly in membrane fractions, and β-actin, a cytoskeleton protein, was found mostly in cytoplasmic fractions (Fig. 3E).

The fractionation of B cells produced distinct luminescence results depending on whether EVs were added to whole blood or purified B cells. In whole blood samples, the luciferase signal was detected in both compartments but was more abundant in the membrane fraction than in the cytoplasmic fraction (Fig. 3F). Particularly at 30 minutes, cytoplasm fractions showed higher values than at 15 or 60 minutes, corroborating our earlier EGFP quantification results from confocal microscopy images. The luminescence was 10-fold less intense in purified B cells than for B cells in whole blood (Fig. 3G). While most of the luciferase signal was detected in the membrane, it was almost undetected in the cytoplasm. These results suggest that, although B cell internalization of EVs can be detected using both experimental designs, external components in whole blood may enhance association and promote distinct internalization mechanisms.

## Discussion

The finding that certain EVs interact with NHP B cells as soon as 10 minutes after intravenous EV administration raised several important questions, including whether these results would also apply to human cells (7). An *ex vivo* model successfully replicated the previous *in vitro* study and can now be used to investigate the mechanisms underlying EV tropism for B cells with minimal or no animal experimentation (10). In this paper, we explored the interactions between EVs and B cells, assessing the applicability of our model to human blood samples and examining how these interactions may diverge from *in vitro* assays using purified B cells.

The association of specific EVs with specific cells is highly relevant for both diagnostics and therapeutics. Classic findings from 1996 showed that EVs carry functional peptide-MHC complexes, suggesting a role in B cell antigen presentation, and a recent review compiled updated references that confirmed this hypothesis (21,22). B cells are vital components of the adaptive immune system, classified by their function and relative expression of surface markers into transitional, immature, marginal zone (MZB), and memory cells. Each B cell ontology has specific functions, which can be further heightened by activation (23,24). The tentative role of EVs in the activation of B cells via the expression of a functional B cell receptor (BCR) has been suggested both in human rheumatoid arthritis and in mouse lymphoma studies (23,24).

However, there is a gap in the translation of basic EV research into clinical practice, especially due to our limited knowledge of EV biology in complex organisms. The use of animal models partially solves this issue and is an invaluable asset to biological research. For instance, NHPs are often studied in pre-clinical research due to their physiological similarities with humans (7). Even so, animal model results are not always recapitulated in human subjects, and other challenges of animal research are worth considering (25). Novel models that can faithfully replicate results obtained with *in vivo* models, such as experimentation with blood *ex vivo*, can sometimes offer a biologically complex and compelling alternative to using living organisms in any way except as biofluid donors (10). In this study, an *ex vivo* model revealed that EV uptake mechanisms by B cells appear to function for both NHP and human cells, supporting previous *in vivo* observations (7). Our main limitation was the maximum sample stability time at a physiological temperature, optimized for up to 4 hours after EV spike-in; this did not allow investigation of lengthier processes such as B cell differentiation and activation (24).

Several comprehensive reviews have described the multiple internalization mechanisms of EVs, including clathrin-dependent and -independent processes and membrane fusion (1,21). A recent study described that EV uptake in mammalian cells is a low-yield process, driven primarily by membrane fusion that requires endosomal acidification for luminal cargo delivery (20). In HeLa cells, the uptake of EVs has been reported to be favored by clathrin-independent mechanisms and macropinocytosis, as discovered via pharmacological inhibition of uptake pathways (26). In our system, approximately half of the EV signal was found internalized in B cells after 30 minutes of spike-in, and only a fraction seems to be metabolized by cells after uptake. We also found no single predominant uptake mechanism, suggesting that clathrin-dependent, clathrin-independent, and direct fusion pathways may be involved in this process. Engaging multiple uptake pathways may offer advantages for selectively modulating the uptake by specific B cell subtypes or for influencing downstream processes such as activation or differentiation into effector populations.

Interestingly, spike-in results differed between purified B cells and B cells in whole blood. Possibly, B cells may change during the manipulation of the purification process, rendering them relatively refractory to association with EVs. However, it is also possible that factors in whole blood could promote EV-B cell associations, including uptake. A recent study found that human blood proteins form a “biomolecular corona” around EVs, improving their uptake by monocytes, suggesting that this mechanism may be used to improve EV clearance from the bloodstream (27). Another report discussed how albumin can prolong the half-life of EVs in circulation (28). In our data, EV interactions with other NHP PBMCs, including monocytes, are also suggestive of particle clearance by phagocytic immune cells (29).

Finally, we would like to address some possible future directions. While EV uptake by B cells appears to be evolutionarily conserved between two primate species, the sensitivity of B cell response to particle uptake differed somewhat. Considering that B cell subpopulations can vary considerably in frequency and absolute counts between healthy humans and with factors such as age (30), we posit that differences in B cell populations might account for observed differences in B cell-EV interactions; this could be tested in future studies. EVs have been suggested to modulate B cell activation, particularly in autoimmune conditions like rheumatoid arthritis (31). The association of EVs and cells may also change with particle surface modifications: for instance, EVs containing surface IgM monomers derived from B cells can bind to specific antigens and aid immune modulation (32). These functional roles of EVs in the B cell world deserve additional attention. Lastly, further studies of uptake mechanisms are needed to realize the potential of EVs for therapeutically modulating healthy B cells and treating B cell diseases.

## Author’s contribution

BCP: conceptualization, data curation, formal analysis, investigation, methodology, project administration, writing – original draft; BVR: conceptualization, formal analysis, methodology, visualization; ENS: methodology, formal analysis, writing – review and editing; OG: methodology, writing – review and editing; BC: methodology, visualization; SEQ: methodology, visualization; LDE: conceptualization, methodology, writing - original draft; HSSA: funding acquisition, supervision, writing – review and editing; KWW: conceptualization, funding acquisition, supervision, resources, writing – review and editing.

## Acknowledgments

The authors thank members of the Witwer Laboratory, the Retrovirus Laboratory, and the Biochemistry and Molecular Biology Laboratory for helpful suggestions and discussions.

## Data availability

The data generated in this study are available within the article and its supplementary data files. Raw files are available upon reasonable request to the corresponding author. We have submitted all relevant data of our experiments to the EV-TRACK knowledgebase (EV-TRACK ID: EV250009) (12). This study does not contain artificial intelligence-generated data.

## Ethics statement

The authors state that all research conducted in this study adhered to the highest ethical standards, ensuring integrity in research practices and compliance with institutional and legal requirements.

