## Supplementary material for "An *ex vivo* and *in vitro* investigation of extracellular vesicle interactions with B cells of *Macaca nemestrina* and humans": Suppl.

#### Table of contents

**Supplementary Figure 1 –** Uncut membranes and gels for EV characterization S2

**Supplementary Information 2 –** MIFlowCyt-EV Framework S3

**Supplementary Information 3 –** MIFlowCyt Framework S7

**Supplementary Figure 4 -** Flow cytometry gating strategy for PBMC separation S12

**Supplementary Figure 5 -** Uncut membranes and gels from cell fractionation S13

**Supplementary Figure 6 -** EV characterization S14

**Supplementary Figure 5 –** PBMC association with $EGFP^+$ EVs in primates S15

### Supplementary Figure 1 – Uncut membranes and gels for EV characterization

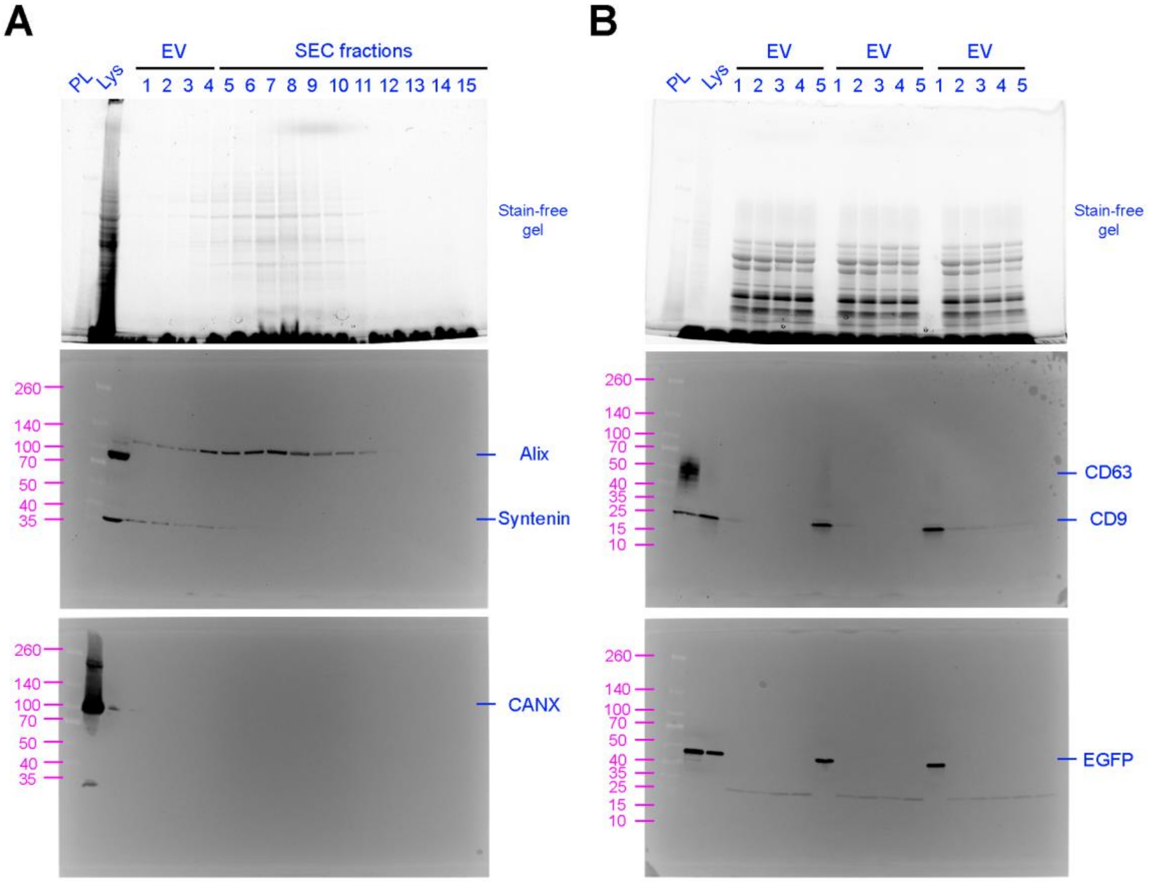


(A) Stain-free gel for map 1 with the screening of SEC fractions from Expi293F-palmGRET EV separation. Membranes probed for Alix, Syntenin and CANX. (B) Stain-free gel for map 2 with the screening of distinct Expi293F-palmGRET EV preparations; EV1 was used in this study. Membranes probed for CD63, CD9 and EGFP. The protein ladder molecular weight is described in magenta.

### Supplementary Information 2: MIFlowCyt-EV framework

| Component | Description |
|---|---|
| **1.1** | EVs were separated from Expi293™ cells (Gibco #A14527) after three days of transfection with the pLenti-palmGRET reporter plasmid (Addgene #158221). EVs were collected by conditioned media centrifugation (300 x g for 5 min, 2,000 x g for 20 min), filtration (0.22 µm bottle-top system, Corning), tangential flow filtration (TFF, Vivaflow® 50R TFF cassettes, Sartorius), ultrafiltration (Centricon Plus 70 Ultracel® PL-100, Merck Millipore; 4,000 x g, 20 min, RT) and size-exclusion chromatography (SEC; qEV10 70nm columns, IZON). From the 11 fractions, EV-enriched fractions 1-4 were pooled and concentrated using Amicon 15 Ultra RC 10 kDa MWCO filters (Merck Millipore). Expi293F-palmGRET EV aliquots in Dulbecco's phosphate buffered saline (DPBS) were stored at 4 ºC for short-term use and at -80 ºC for long-term storage. |
| **1.2** | **1.1 Aim:** To characterize Expi293F-palmGRET EVs regarding their size, concentration and fluorescence following MISEV2023 guidelines. **1.2 Keywords:** EV; extracellular vesicles; characterization. **1.3 Experimental variables:** none. |
| **2.1** | The Expi293™ expression system (Gibco #A14635) was used to transiently transfect the pLenti-palmGRET reporter plasmid. Expi293 cells were seeded in sterile Erlenmeyer flasks in Expi293 growth media under a 3 x $10^6$ cells/ml concentration and allowed to grow overnight at 37 ºC, 8% $CO_2$ under orbital shaker. Cells were diluted to a 3 x $10^6$ cells/ml concentration in fresh media. Two solutions containing the ExpiFectamine™ 293 reagent or the pLenti-palmGRET plasmid (1 µg total plasmid DNA per mL of culture volume) in Opti-MEM™ I Medium were prepared and incubated for 5 minutes at room temperature. Both solutions were combined at room temperature for 10 min before being added to the cell culture while swirling. Cells were maintained at 37 ºC, 8% $CO_2$ for 18-22 hours, and checked for EGFP fluorescence using a microscope. A transfection enhancer was added to the cell culture, and the flasks were further incubated for two days. On day 3 after transfection, $EGFP^+$ EVs were collected from the conditioned media. |
| **2.2** | Cells were harvested from flasks and centrifuged at 300 x g for 5 min to separate from the conditioned media. The media was further centrifuged at 2,000 x g for 20 min to remove debris and filtered using a bottle-top system (0.22 µm). After tangential flow filtration and ultrafiltration, the conditioned media was separated using size-exclusion chromatography using DPBS as buffer. |
| **2.3** | 1 µl of EV samples were diluted in 999 µl of DPBS for a 1:1000 dilution. For concentrated samples, a 100 µl aliquot of the 1:1000 dilution was further diluted in 900 µl of DPBS, resulting in a 1:10,000 dilution. |
| **3.1** | We used as buffer-only control a 0.22 µm filtered DPBS sample, which was recorded prior to the analysis using the same acquisition parameters. The buffer-only control had a count of ~0.98 events per second. |
| **3.2** | No buffer with reagents were used in this experiment. |
| **3.3** | No unstained controls were used in this experiment. |
| **3.4** | No isotope controls were used in this experiment. |
| **3.5** | No single-stained controls were used in this experiment. |
| **3.6** | No procedural controls were used in this experiment. |
| **3.7** | Samples were serially diluted for up to two times, with 1 µl of EV samples diluted in 999 µl of DPBS for a 1:1000 dilution. If samples were too concentrated, a 100 µl aliquot of this 1:1000 dilution was further diluted in 900 µl of DPBS, resulting in a 1:10,000 |
| **3.8** | No detergent-treated controls were used in this experiment. |

| Component | Description |
|---|---|
| **4.1** | Based on the buffer alone control (**Component 3.1**), detection was triggered on the 488 nm laser excited FITC channel (10-50 mW 488 laser) at a threshold of 13.7 arbitrary units, determined using quality control beads (0.25 µm Fluorescent Silica Microspheres) and the manufacturer's calibration values (see **Component 4.3** for fluorescence calibration). The buffer alone control had an event rate of ~1 events per second. |
| **4.2** | Samples were enumerated using the integral instrument flow rate sensors, resulting in a flow rate of 26.07 nL/min. This was calibrated using weighted volumes of deionized water prior to analysis. |
| **4.3** | Arbitrary FITC fluorescence scale units (channel number), excited by the 488 nm laser and collected using a 525/40 bandpass filter were converted to MESF units using 0.25 µm Fluorescent Silica Microspheres (250nm Std Fi SiNPs, NanoFCM Inc). |
| **4.4** | Side scatter calibration was performed on the NanoAnalyzer software (NanoFCM Inc), considering the wavelength (405 nm) and polarization state (perpendicular to detection of the laser), the light collection geometry (side scatter, numerical aperture) and the particle diameter and refractive index. Side scatter decay was reported at 10%. |
| **5.1** | EV diameter was approximated using a calibration curve determined by silica nanospheres size and side scattering intensity. The silica nanosphere cocktail contained particles with diameters ranging from 68~155 nm (S16M-Exo, NanoFCM Inc), peaking at 68, 91, 113 and 155 nm. Beads were boosted under 10 mW laser power, 10% SS decay and ≤1.0 kPa sampling pressure. |
| **5.2** | Particle refractive index was derived from the ratio of side and forward scatter signal following the silica nanosphere calibration curve described in **Component 5.1**. |
| **5.3** | No EV epitopes were evaluated in this experiment. |
| **6.1** | The MIFlowCyt checklist v.1.0.0 has been completed and attached below. |
| **6.2** | The calibrated FITC channel detected nanoparticles ranging from 40-200 nm diameters and up to 407,000 FITC MESF units. Positive events on FITC were assumed to be higher than a threshold of 57.8 FITC MESF units. |
| **6.3** | The detected concentration of Expi293F-palmGRET EVs between calibrated detection range (57.8 and 407,000 FITC MESF units) was 4.65 x $10^{11}$ particles per ml. Flow cytometer acquisition settings were maintained for all samples, including triggering threshold, voltages and flow rate. |
| **6.4** | EV brightness was determined as a 78.7% fraction of the total event count, corresponding 9906 of the 12582 events recorded. EGFP brightness in EV samples was not quantified. |
| **7.1** | NFA files and reports are available upon reasonable request by contacting the corresponding author. |

### MIFlowCyt – EV characterization

| Component | Description |
|---|---|
| **1. Experimental Overview** | |
| **1.1 Purpose** | To characterize Expi293F-palmGRET EVs regarding their size, concentration and fluorescence following MISEV2023 guidelines |
| **1.2 Keywords** | EV; extracellular vesicles; characterization. |

| Component | Description |
|---|---|
| **1.3 Experimental Variables** | None. |
| **1.4 Organization** | ***1.4.1 Name:*** The Retrovirus Laboratory, Department of Molecular and Comparative Pathobiology, Johns Hopkins University School of Medicine<br>***1.4.2 Address:*** 733 North Broadway, Miller Research Building, Baltimore, MD 21205, United States |
| **1.5 Primary Contact** | ***1.5.1 Name:*** Dr. Kenneth Whitaker Witwer<br>***1.5.2 Email Address:*** |
| **1.6 Date** | Flow cytometry analysis was performed on Feb 2, 2023. |
| **1.7 Conclusions** | We obtained an EV preparation with a concentration of 4.65 x $10^{11}$ particles per ml and 78.7% of particles were positive for EGFP fluorescence. |
| **1.8 Quality Control Measures** | The system was calibrated using quality control beads and a vehicle control (i.e., DPBS) was used as a blank. |

### 2. Flow Sample/Specimen Details

#### 2.1 Sample/Specimen Material Description

| | |
|---|---|
| ***2.1.1 Biological Samples*** | *2.1.1.1 Biological Sample Description:* EVs separated from the conditioned media from Expi293F human cells (Gibco # A14527) transfected using the Expi293F™ expression system (Gibco # A14635) with the plasmid pLenti-palmGRET (Addgene #158221).<br>*2.1.1.2 Biological Sample Source Description:* The Expi293F cells are derived from the HEK 293 human embryonic kidney cell line and are grown under suspension in Expi293™ Expression Medium (Gibco #A1435101).<br>*2.1.1.3 Biological Sample Source Organism Description:* EVs separated from the conditioned media from Expi293F human cells transfected using the Expi293F™ expression system with the plasmid pLenti-palmGRET.<br><u>2.1.1.3.1 Taxonomy:</u> *Homo sapiens*<br><u>2.1.1.3.4: Phenotype:</u> Expi293F human cells are derived from the 293 cell line, and are a core component of the Expi293 Expression System. They are maintained in suspension culture and will grow to high density in Expi293 Expression Medium. Expi293F cells are highly transfectable and generate superior protein yields compared to standard 293 cell lines in transient protein expression.<br><u>2.1.1.3.5: Genotype:</u> Undisclosed by the manufacturer (Thermo Fisher Scientific).<br><u>2.1.1.3.6 Treatment:</u> The Expi293™ expression system (Gibco #A14635) was used to transiently transfect the pLenti-palmGRET reporter plasmid. Expi293 cells were seeded in sterile Erlenmeyer flasks in Expi293 growth media under a 3 x $10^6$ cells/ml concentration and allowed to grow overnight at 37 ºC, 8% $CO_2$ under orbital shaker. Cells were diluted to a 3 x $10^6$ cells/ml concentration in fresh media. Two solutions containing the ExpiFectamine™ 293 reagent or the pLenti-palmGRET plasmid (1 µg total plasmid DNA per mL of culture volume) in Opti-MEM™ I Medium were prepared and incubated for 5 minutes at room temperature. Both solutions were combined at room temperature for 10 min before being added to the cell culture while swirling. Cells were maintained at 37 ºC, 8% $CO_2$ for 18-22 hours, and checked for EGFP fluorescence using a microscope. A transfection enhancer was added to the cell culture, and the flasks were further incubated for two days. On day 3 after transfection, $EGFP^+$ EVs were collected from the conditioned media. |

| Component | Description |
|---|---|
| **2.3 Sample Treatment(s) Description** | None |
| **2.4 Fluorescence Reagent(s) Description** | EV samples were separated after transfection with the EGFP tag and detected accordingly to the following optical detectors:<br>- *Reporter for FL1:* SS<br>- *Reporter for FL2:* FITC<br>- *Reporter for FL3:* PC5 |
| **3 Instrument Details** | |
| **3.1 Instrument Manufacturer** | NanoFCM Inc<br>https://www.nanofcm.com |
| **3.2 Instrument Model** | Flow NanoAnalyzer<br>https://www.nanofcm.com/category/productorcenter/flownanoanalyzer/ |
| **3.3 Instrument Configuration and Settings** | Instrument configuration and settings were consistent with the original installation standard (June 28, 2018). |

### Supplementary Information 3 – MIFlowCyt framework

| Component | Description |
|---|---|
| **1. Experimental Overview** | |
| **1.1 Purpose** | The experiment aimed to phenotypically separate peripheral blood mononuclear cells, including $CD20^{+}$ B cells, in the whole blood of macaques and humans spiked with Expi293F-palmGRET EVs *ex vivo*. |
| **1.2 Keywords** | Extracellular vesicles; B cells; *Macaca nemestrina*; Human; Flow cytometry |
| **1.3 Experimental Variables** | We wanted to detect the uptake of $EGFP^{+}$ EVs by cells from the PBMC pool of macaques and human samples. Hence, samples were evaluated by flow cytometry, mixed with trypan blue, and re-analyzed. |
| **1.4 Organization** | ***1.4.1 Name****:* The Retrovirus Laboratory, Department of Molecular and Comparative Pathobiology, Johns Hopkins University School of Medicine<br>***1.4.2 Address:*** 733 North Broadway, Miller Research Building, Baltimore, MD 21205, United States |
| **1.5 Primary Contact** | ***1.5.1 Name:*** Dr. Kenneth Whitaker Witwer<br>***1.5.2 Email Address:*** |
| **1.6 Date** | Blood draws on human and pigtailed macaque samples included in this experiment range from September 2022 to August 2023. Analyses ranged from September 2022 through August 2023. |
| **1.7 Conclusions** | In NHP PBMCs, the EGFP signal was detected after 10 min of EV spike-in in B cells (72.53% ± 9.57%), $CD14^{+}$ monocytes (89.6% ± 7.15%), $CD14^{-/lo}$ monocytes (63.68% ± 10.59%), CD4 T lymphocytes (19.63% ± 11.90), CD8 T lymphocytes (6.62% ± 2.76%). Little or no interaction of EVs with CD159a+ NK cells was detected (0.05% ± 0.08%). In human PBMCs, the EGFP signal was present in 65.33%, 71.35%, and 81.70% of B cells after 15, 30, and 60 min. Internal EGFP signal in human B cells was present in 27.52%, 37.72% and 55.33% of the population after 15, 30 and 60 min, respectively. |
| **1.8 Quality Control Measures** | For samples acquired on different days, voltage settings were standardized to daily CS&T Research Bead (BD Biosciences, San Jose, CA) controls using predetermined application settings in FACSDiva 9.0 to ensure fluorescence intensity was consistent longitudinally. An FMO control was used for gating of GFP samples in each experiment. |
| **2. Flow Sample/Specimen Details** | |
| **2.1 Sample/Specimen Material Description** | |
| ***2.1.1 Biological Samples*** | *2.1.1.1 Biological Sample Description:* Human whole blood was collected intravenously using citrate-phosphate-dextrose (CPD) blood bags. Pigtailed macaque whole blood was collected intravenously using acid citrate dextrose (ACD) syringes.<br>*2.1.1.2 Biological Sample Source Description:* Human whole blood or pigtail macaque whole blood.<br>*2.1.1.3 Biological Sample Source Organism Description:* Human whole blood bags were purchased from the New York Blood Center (New York, NY, USA) for research use only. Macaque whole blood was collected from healthy pigtails.<br>2.1.1.3.1 Taxonomy: *Homo sapiens*; *Macaca nemestrina* |

| Component | Description |
|---|---|
| | 2.1.1.3.2 Age: For human donors, ages ranged from 29 to 67 years old at time of donation. For pigtail macaques, blood was drawn from juvenile and adult individuals.<br><br>2.1.1.3.3 Gender: Six human female donors, one human male donor, two female macaque donors and five male macaque donors.<br><br>2.1.1.3.4: Phenotype: Healthy individuals only.<br><br>2.1.1.3.6: Treatment: Within one hour (for macaque) and one day (for human) of collection, whole blood was aliquoted into conical tubes and mixed with Expi293F-palmGRET EVs (8E+8 particles/ml). Samples were incubated on a rotator in an incubator set at 37 ºC, with the "mix" mode selected at a speed of 8 rpm (Benchmark Scientific Roto-Therm Plus Incubated Rotator). EV interaction with whole blood components was monitored at 5, 10, 15, 30, 60, 120, and/or 240 minutes after spike-in.<br><br>2.1.1.3.7. Other Relevant Biological Sample Source Organism Information: Animal studies were approved by the Johns Hopkins Institutional Animal Care and Use Committee and conducted in accordance with the Weatherall Report, the NIH Guide for the Care and Use of Laboratory Animals, and the USDA Animal Welfare Act. Animals were monitored twice daily by veterinarians or trained technicians for signs of clinical disease. Macaques were housed in Johns Hopkins University facilities that are fully accredited by the Association for the Assessment and Accreditation of Laboratory Animal Care, International (AAALAC), and fed a balanced commercial macaques chow (Purina Mills, St. Louis, MO). All animals received environmental enrichment including manipulanda and novel foodstuffs throughout the study and were pair-housed. |
| ***2.1.2 Environmental Samples*** | None |
| ***2.1.3 Other Samples*** | BD CompBead Anti-mouse Ig, κ/Negative particles (BD Biosciences, San Jose, CA). |
| **2.2 Sample Characteristics** | All cell types included in whole blood, with the exception of red blood cells, which are lysed during RBC lysis treatment, are included in analyses. |
| **2.3 Sample Treatment(s) Description** | Within one hour (for macaque) and one day (for human) of collection, whole blood was aliquoted into conical tubes and mixed with Expi293F-palmGRET EVs (8E+8 particles/ml). Samples were incubated on a rotator in an incubator set at 37 ºC, with the "mix" mode selected at a speed of 8 rpm (Benchmark Scientific Roto-Therm Plus Incubated Rotator). EV interaction with whole blood components was monitored at 5, 10, 15, 30, 60, 120, and/or 240 minutes after spike-in.<br><br>NHP blood samples were diluted (1:1, v/v) in Hank's balanced salt solution (HBSS) and applied to SepMate™-15 (IVD) tubes loaded with Percoll® density gradient medium (Cytiva #17-0891-01). Tubes were centrifuged at 1,200 × g for 10 min for triphasic blood separation, with mononuclear cells (MNC) in the upper fraction. MNCs were transferred to fresh tubes, washed with HBSS, and pelleted at 300 × g for 8 min. Cells were treated with red blood cell lysis buffer (ACK lysing buffer 0.83% $NH_4Cl$, 0.1% $KHCO_3$, 0.03% EDTA) for 10 min at 37 ºC, washed with HBSS, and centrifuged at 300 × g for 8 min.<br><br>Human donor (HD) blood samples were diluted (1:1, v/v) in washing buffer (PBS + 2% FBS + 2 mM EDTA) and applied to SepMate™-50 (IVD) tubes loaded with Ficoll-Paque™ PLUS density gradient (Cytiva # 17144003). Tubes were centrifuged at 1,200 × g for 10 min for triphasic blood separation, with mononuclear cells (MNC) in the upper fraction. MNCs were transferred to fresh tubes, washed with buffer, and pelleted at 300 × g for 8 min. Cells were treated with red blood cell lysis buffer for 10 minutes at 37 ºC, |

| Component | Description |
|---|---|
| | washed with buffer, and centrifuged at 300 × g for 8 min. MNCs were stained with a pre-titrated amount of monoclonal antibodies in a dark hood at room temperature for 20 minutes. Labeled PBMCs were washed once with DPBS (300 × g, 8 min), resuspended in DPBS, and aliquoted (500 µl) for PBMC-associated EGFP fluorescence analysis. |

**2.4 Fluorescence Reagent(s) Description**

| Analyte | Analyte Detector | Analyte Reporter | Reagent Manufacturer | Reagent Catalogue |
|---|---|---|---|---|
| CD159a | Anti-CD159a antibody | PE | Beckman Coulter | IM3291U |
| CD4 | Anti-CD4 antibody | PerCP-Cy5.5 | BD Biosciences | 552838 |
| CD20 | Anti-CD20 antibody | eFluor™ 450 | Invitrogen | 48-0209-42 |
| CD3 | Anti-CD3 antibody | V500 | BD Biosciences | 560770 |
| CD8a | Anti-CD8a antibody | Brilliant Violet 570 | BioLegend | 301038 |
| CD14 | Anti-CD14 antibody | Brilliant Violet 650 | BD Biosciences | 563419 |
| EVs | - | GFP | - | - |

### 3 Instrument Details

**3.1 Instrument Manufacturer**

BD Biosciences, San Jose, CA

https://www.bdbiosciences.com/en-us

**3.2 Instrument Model**

BD LSRFortessa

**3.3 Instrument Configuration and Settings**

Software Version: BD FACSDiva 9.0. Configuration Name Fortessa 6-Blue 6-Violet 3-Red. Operational system: Window Extension 10.

***3.3.4 Optical Filters***

See table below. Optical Filters are original to manufacture date of July 2010.

***3.3.5 Optical Detectors***

See table below.

| Laser Name | Type | Wavelength | Power | Detector Array | Detector | Channel | Mirror (LP) | Filter (BP) | Parameter | Position | Voltage |
|---|---|---|---|---|---|---|---|---|---|---|---|
| Blue | Blue | 488 | 20 | Octagon | A | 6 | 750 | 780/60 | PE-Cy7 | 1 | - |
| | | | | | B | 5 | 685 | 695/40 | Per-CP-Cy5.5 | | 663 |
| | | | | | C | 4 | 600 | 610/20 | PE-Texas Red | | - |
| | | | | | D | 3 | 550 | 575/26 | PE | | 537 |
| | | | | | E | 2 | 505 | 530/30 | Alexa Fluor 488 | | 544 |
| | | | | | F | 1 | | | | | |
| Violet | Violet | 405 | 25 | Octagon | A | 12 | 630 | 655/8 | Brilliant Violet 650 | 2 | 685 |
| | | | | | B | 11 | 595 | 605/12 | Brilliant Violet 605 | | - |
| | | | | | C | 10 | 575 | 585/15 | Qdot 585 | | - |
| | | | | | D | 9 | 545 | 560/20 | Qdot 565 | | 626 |
| | | | | | | | | | Brilliant Violet 570 | | |
| | | | | | E | 8 | 475 | 525/50 | AmCyan | | 602 |
| | | | | | | | | | V500 | | |
| | | | | | | | | | Qdot 525 | | |
| | | | | | F | 7 | | 450/50 | Pacific Blue | | 526 |
| | | | | | | | | | Brilliant Violet 421 | | |
| | | | | | | | | | VioBlue | | |
| | | | | | G | | | | | | |
| | | | | | H | | | | | | |
| Red | Red | 633 | 17 | Trigon | A | 15 | 755 | 780/60 | APC-Cy7 | 3 | - |
| | | | | | B | 14 | 710 | 730/45 | Alexa Fluor 700 | | 616 |
| | | | | | C | 13 | | 670/14 | APC | | - |

| Component | Description |
|---|---|
| **4. Data Analysis Details** | |
| **4.1 List-mode data files** | FCS data files can be obtained upon request to the corresponding author. |
| **4.2 Compensation Details** | Data were compensated in BD FACSDiva 9.0 software (BD Biosciences, San Jose, CA) using the Compensation feature. Anti-mouse Ig, κ/Negative Control CompBead (BD Biosciences, San Jose, CA) were antibody stained with matched fluorochrome or matched lot and tube in the case of Brilliant Violet dyes, and a universal control negative lot-matched bead. CD3 V500 single-color stained whole blood was used and negative whole blood gated for the V500 compensation particle. GFP positive and negative samples were used for GFP compensation. The compensation matrix can be obtained upon request to the corresponding author. |
| **4.3 Data transformation details** | ***4.3.1 Purpose of Data Transformation***: Data was transformed to enhance visualization of all fluorochromes on scale.<br>***4.3.2 Data Transformation Description:*** Biexponential transformation of fluorochrome data was utilized in FlowJo 10.10.0 software (BD, Ashland, OR). |
| **4.4 Gating (Data Filtering) Details** | ***4.4.1 Gate Description:*** Events were first gated using FSC-A vs. FSC-H to remove doublets. Cells were then gated based on size and complexity using FSC-A vs. SSC-A to choose events of relative particle shape and size of interest to remove debris. Those cells were then gated as lymphocytes or monocytes further based on FSC-A and SSC-A characteristics. Lymphocytes were then gated as CD3+ CD20- T cells, CD3- CD20+ B cells, or CD3- CD20- lymphocytes. CD3+ T cells were further gated as CD4+ CD8- T helper cells or as CD8+ CD4- cytotoxic T lymphocytes. CD3- CD20- lymphocytes were then gated as CD159a+ NK cells. Monocytes were further refined as CD159a-, CD3-, CD20- monocytes. Those monocytes were then gated for CD14 expression as CD14+ or CD14 low. Each cell subset was gated for EGFP positivity with EGFP expression based on no EGFP controls. Gating is shown in Supplementary Figure 2 using FlowJo 10.10.0 software (BD, Ashland, OR).<br>***4.4.2 Gate Statistics:*** Quantitative data for the proportion of B lymphocytes expressing EGFP shown in Figures 1E, 1H, and 3A used the gating strategy shown below in Supplementary Figure 4. In detail, the B cells were characterized as the singlet→cell discrimination from FSC-A vs. SSC-A→lymphocyte discrimination from FSC-A vs. SSC-A→CD20+ CD3- B lymphocytes. The CD20+ CD3- B lymphocytes are then gated as percentage of cells expressing EGFP as based on the no EGFP control shown in Figure 4. Supplementary Figure 7 uses the same gating strategy shown in Supplementary Figure 4. Each additional lymphocyte and monocyte subtype is then gated for EGFP expression based on the no-EGFP control (not shown). Gates depicted in all figures were calculated using FlowJo 10.10.0 software (BD, Ashland, OR).<br>***4.4.3 Gate Boundaries:*** Gate boundaries have been shown for all figures included in the paper, with the exception of the EGFP boundaries for the cell subsets only included in Supplementary Figure 7, in Supplementary Figure 4. |

**Supplementary Figure 4 – Flow cytometry gating strategy for PBMC separation**

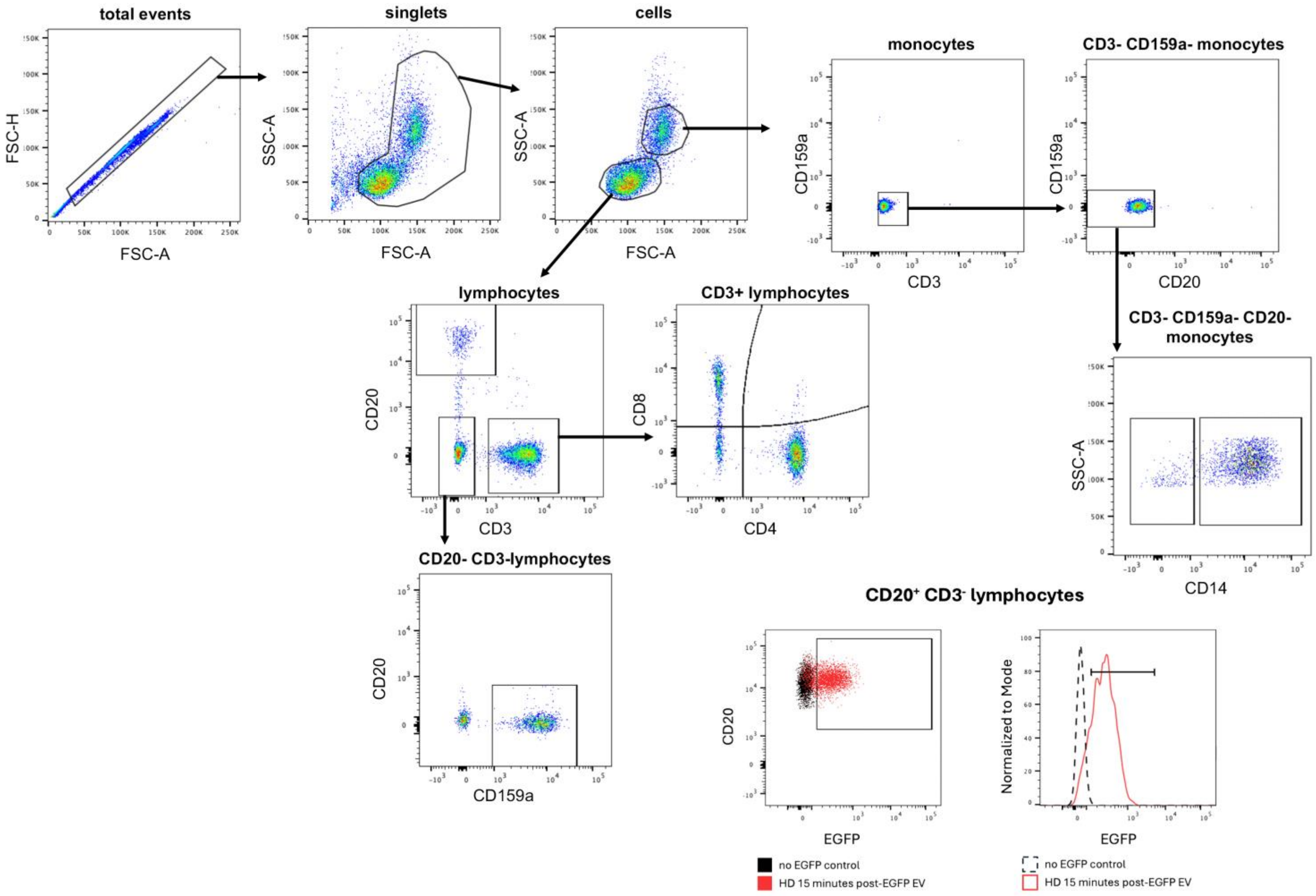


Gating and analysis were produced on the FlowJo 10.10.0 software (BD, Ashland, OR).

**Supplementary Figure 5 - Uncut membranes and gels from cell fractionation**

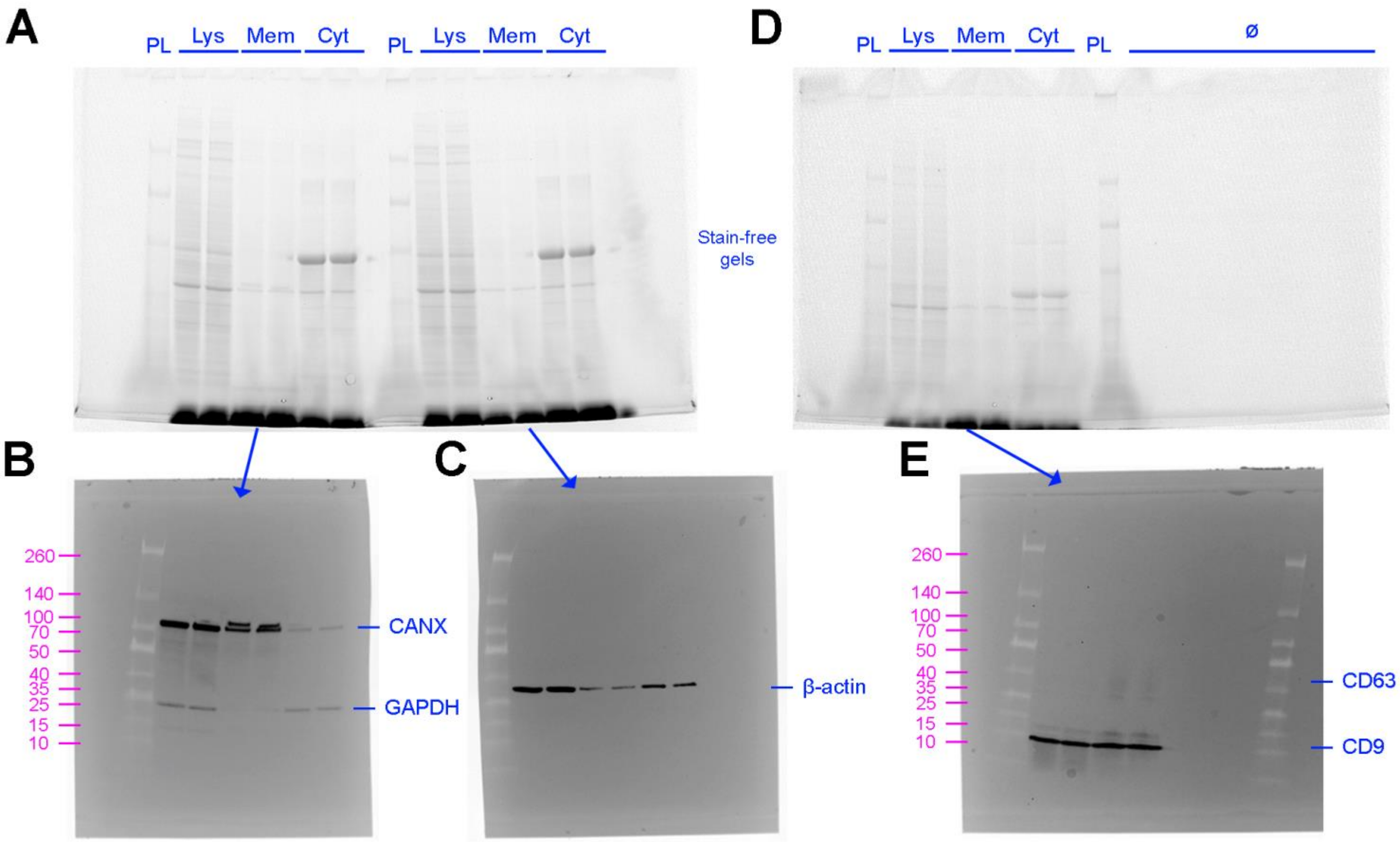


(A) Stain-free reducing gel loaded with a protein ladder (PL), full cell lysate (Lys), membrane fractions (Mem) and cytoplasmic fractions (Cyt) of human purified B cells. (B-C) Membranes from the reducing gel were cut to probe for (B) calnexin (CANX) and GAPDH, or (C) β-actin. (D) Stain-free non-reducing gel map, with extra wells left empty. (E) Membrane from the non-reducing gel was probed for CD63 and CD9. The protein ladder molecular weight is described in magenta.

### Supplementary Figure 6 - EV characterization

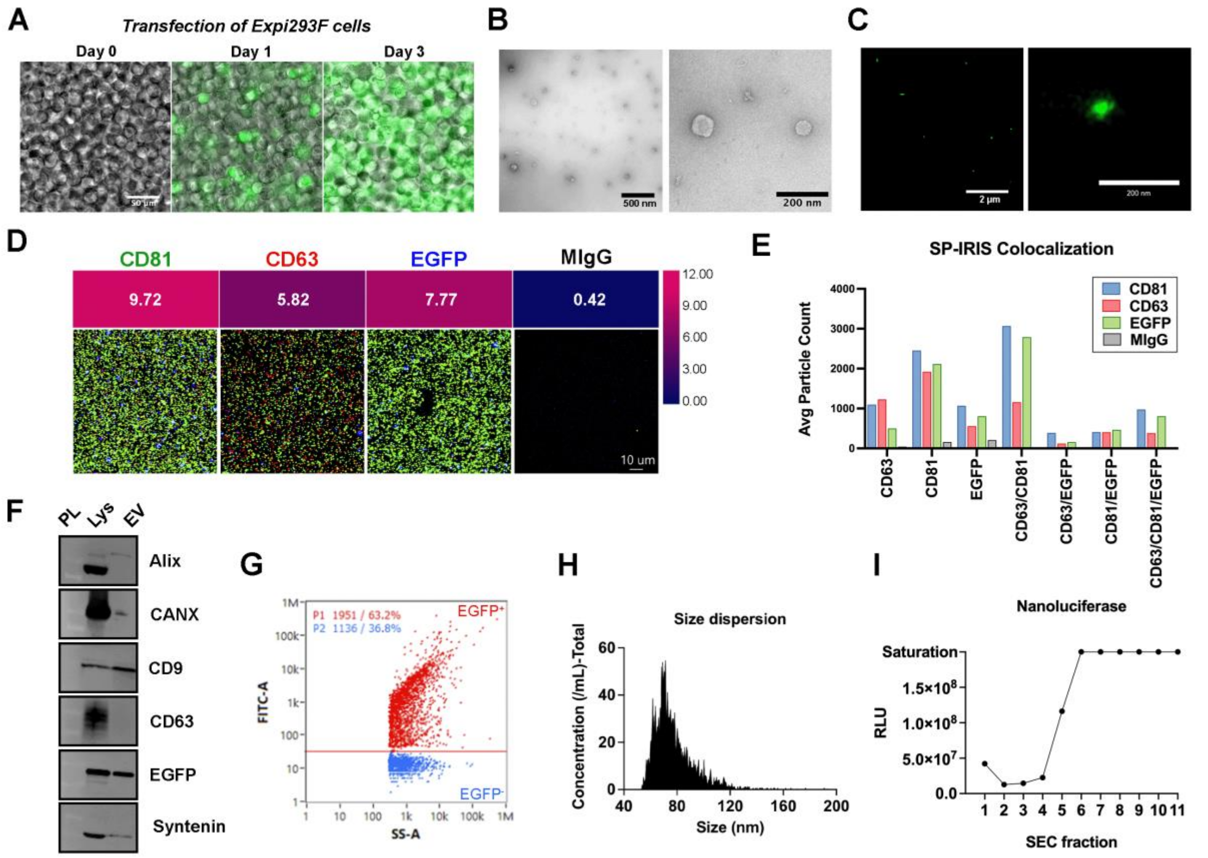


(A) Tracking of Expi293F cell transfection with pLenti-palmGRET by fluorescence microscopy. Scale bar: 50 µm. (B) Widefield and zoomed transmission electron microscopy (TEM) images of Expi293F-palmGRET EVs. Scale bars: 500 nm (left) and 200 nm (right). (C) Widefield and zoomed super-resolution microscopy (SRM) images of EVs separated from Expi293F-palmGRET. Scale bar: 2 µm (left) and 200 nm (right). (D) Co-localization of CD81, CD63, and EGFP in EVs by SP-IRIS. Scale bar: 10 µm. (E) Co-localization plot of average particle count per marker from SP-IRIS. (F) Qualitative western blotting of Alix, calnexin (CANX), CD9, CD63, EGFP, and syntenin in cell lysates and EVs. (G) Scatter plot of EGFP+ EVs (red) obtained from nanoflow cytometry. (H) Diameter distribution plot of EGFP+ EVs, peaking at 79 nm. (I) Relative luminescence units (RLU) of SEC fractions probed by nanoluciferase assay.

### Supplementary Figure 7 – PBMC association with EGFP$^+$ EVs in primates

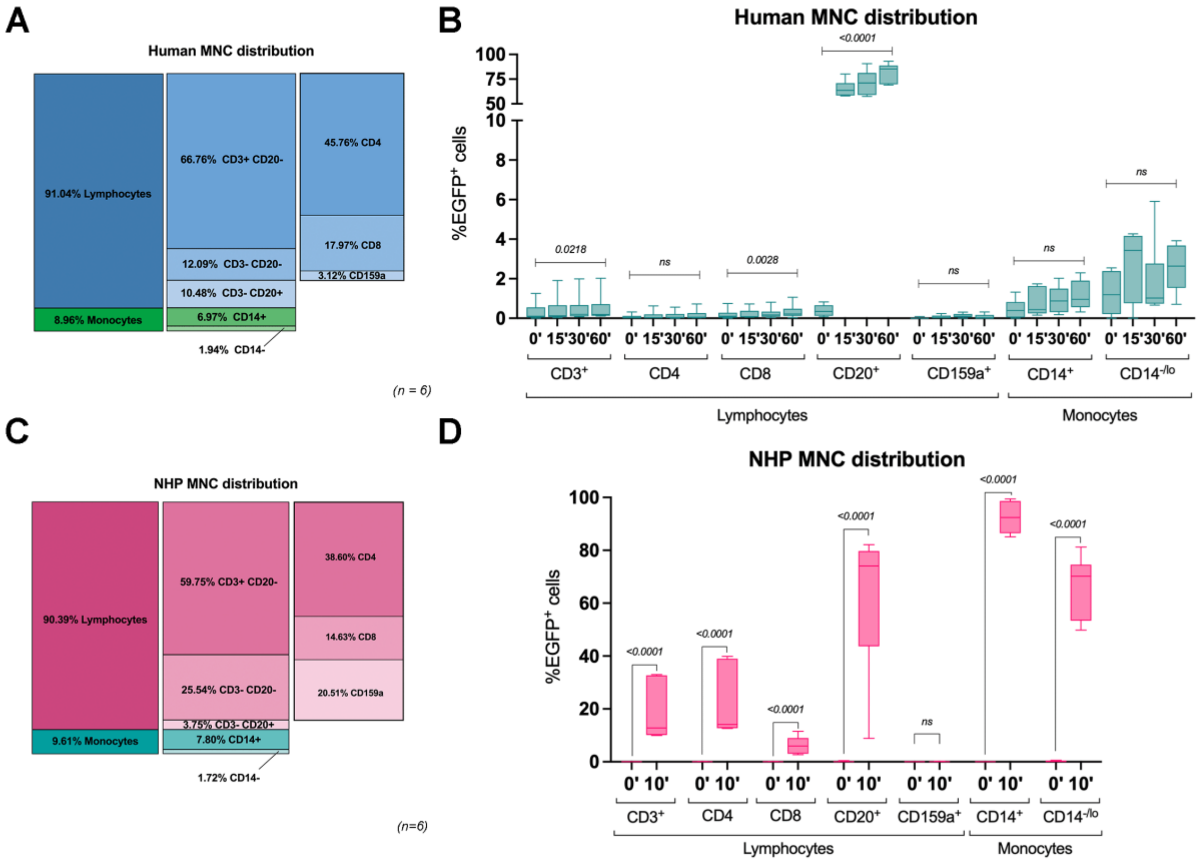


(A) Flow cytometry detected the ratio of PBMC subtypes in human whole blood. Values are presented as the mean of multiple experiments (n=6). (B) Detection of % EGFP signal in human PBMCs after 15, 30, and 60 minutes of Expi293F-palmGRET EV spike-in *ex vivo*. The Friedman test with Dunn's multiple comparisons evaluated the statistical comparison between the values for each cell type, and significant *p* values are displayed above brackets. (C) Ratio of PBMC subtypes in NHP whole blood as detected by flow cytometry. Values are presented as the mean of multiple experiments (n=6). (D) Detection of % EGFP signal in peripheral blood mononuclear cells (PBMC) from non-human primates (NHP) after 10 minutes of Expi293F-palmGRET EV spike-in. The Wilcoxon matched-pairs signed rank test evaluated the statistical comparison between the 0' and 10' values for each cell type, and significant *p* values are displayed above brackets. ns = non-significant.
